# In vivo longitudinal mapping of brain iron accumulation after pilocarpine-induced status epilepticus

**DOI:** 10.64898/2026.03.18.712677

**Authors:** Franco Moscovicz, Leonardo Vazquez-Morales, Alberto Lazarowski, Luis Concha, Jeronimo Auzmendi, Hiram Luna-Munguia

## Abstract

Ferroptosis is a form of non-apoptotic cell death in which iron catalyzes the formation of reactive oxygen species, leading to lipid peroxidation. Experimentally, this process has recently been associated with seizures based on the increased levels of specific markers (4-hydroxynonenal and malondialdehyde) in the brain and plasma. Clinically, iron deposits have been identified in resected tissue from patients with refractory temporal lobe epilepsy. Quantitative susceptibility mapping (QSM) offers an opportunity to detect these accumulations *in vivo*. In this study, we investigated how pilocarpine-induced *status epilepticus* contributes to the generation of iron deposits in diverse cerebral regions and whether QSM can detect these deposits longitudinally. We scanned 14 animals (n=10 experimental; n=4 control) at five different time points (pre-*status epilepticus* induction and 1, 7, 14, 21 days post-induction) using QSM. We identified iron deposits in the caudate putamen, hippocampus, thalamus, and primary somatosensory cortex of experimental animals, which is consistent with histological findings. The initial size of the hippocampal iron deposits significantly increased over the following weeks. None of these effects was observed in the control animals. The presence of cerebral iron depositions in an animal model of pilocarpine-induced *status epilepticus* suggests that ferroptosis may be involved in the onset, development, and progression of spontaneous recurrent seizures. Furthermore, non-invasive, longitudinal *in vivo* mapping of brain iron deposits could be a potential imaging marker in neurological disorders such as epilepsy. Future experiments will be required to determine the origin of the iron and avoid its progressive accumulation.

**Graphical Abstract:** 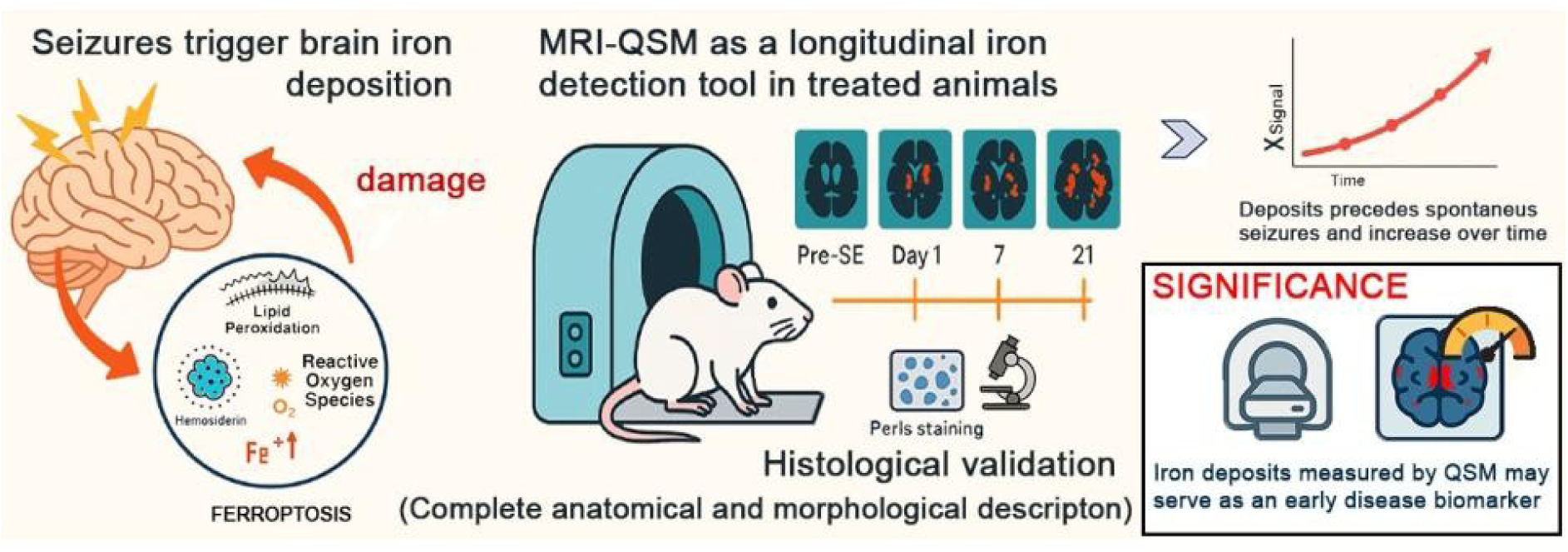

## 1. INTRODUCTION

The biological relevance of iron lies in its capacity to donate electrons, playing a crucial role in diverse cellular processes such as DNA replication, the electron transport chain, and energy production. However, the reaction between iron and molecular oxygen can generate toxic compounds, known as reactive oxygen species (ROS) (Crichton, 2016). The term ferroptosis was coined in 2012 to describe a form of non-apoptotic cell death in which iron, via the Fenton reaction, catalyzes ROS and promotes membrane damage as a consequence of massive lipid peroxidation (Dixon et al., 2012). In contrast to apoptosis and necrosis, ferroptosis has its own morphological, biochemical, and genetic features. These are mainly characterized by mild chromatin condensation, cytoplasm and cytosolic organelle swelling, plasma membrane integrity deterioration, and mitochondrial alterations (Dixon et al., 2012). Diverse studies have also reported that tissues undergoing ferroptosis show increased levels of specific biomarkers, including 4-hydroxynonenal (4-HNE) and malondialdehyde (MDA) (Chen et al., 2023a; Su et al., 2019; Tang et al., 2021; Zhang et al., 2023). Moreover, other works have demonstrated that stress-induced iron accumulations can lead to the generation of hemosiderin, a water-insoluble iron storage protein detectable by Perls staining (Akyuz et al., 2021; Sonoda et al., 2025).

The abnormal accumulation of iron in the brain can trigger inflammation, neurotransmitter oxidation, neuronal communication failure, myelin sheath degeneration, astrocyte dysregulation, and cell death (Foley et al., 2022). These features have been observed in diverse neurological disorders or neurodegenerative processes such as stroke (Alim et al., 2019), traumatic brain injury (Dong et al., 2023), neuroinflammation (Zhang et al., 2022), Alzheimer’s disease (Pomilio et al., 2023), Parkinson’s disease (Do Van et al., 2016), and Huntington’s disease (Tan et al., 2021). In the case of hemosiderin, its cerebral accumulation has been reported in subarachnoid haemorrhage (Hirano et al., 2019; Ikawa et al., 2021), traumatic brain injury (Schuler and Kvistad, 2017), high-altitude cerebral edema (Schommer et al., 2013), and microbleeds secondary to recurrent ischemic strokes or *status epilepticus* (Jeon et al., 2013; Lim et al., 2015). Interestingly, high chances of developing epilepsy have been reported in these cases. Moreover, hemosiderin deposits are also found surrounding brain tumors or cerebral cavernous malformations (Roelcke et al., 2013; Zyck and Gould, 2024); therefore, an incorrect or partial removal may induce seizures (Ruan et al., 2015; Shivamurthy et al., 2024).

During the last five years, researchers have suggested that an imbalanced iron metabolism can contribute to ferroptosis, a process possibly involved in the pathophysiology of epilepsy (Auzmendi and Lazarowski, 2025; Cai and Yang, 2021). The altered antioxidant systems will induce an increase in the expression of the x_c_ ^-^ system (Dahlmanns et al., 2023; Loewen et al., 2019; Martinc et al., 2024; Sarecka-Hujar et al., 2022) and a decrease in the concentration of glutathione and glutathione peroxidase. Experimental studies have also reported increased levels of specific biomarkers (4-HNE, MDA, and cyclooxygenase-2 [COX-2]) during epilepsy development (Folbergrova et al., 2023; Rawat et al., 2022; Sattar et al., 2025). Others have described that inhibiting ferroptosis could attenuate seizures in mice (Chen et al., 2022; Mao et al., 2019) or even exert neuroprotective effects (Shi et al., 2024a). Despite the recent evidence, the dynamics of cerebral iron deposition under convulsive conditions has not yet been fully elucidated.

Quantitative susceptibility mapping (QSM), an advanced magnetic resonance imaging technique, has provided non-invasive insight into tissue susceptibility to iron content variations (Aggarwal et al., 2018; Haacke et al., 2015; Harada et al., 2022). Aggarwal and collaborators (2018) were the first to describe iron accumulations in specific brain regions after *status epilepticus* induction by using pilocarpine-treated rats. Later clinical or experimental imaging findings were further validated by histological or immunohistochemical techniques, confirming the involvement of ferroptosis in epilepsy (Zimmer et al., 2021). The aim of this study was to longitudinally describe the dynamics of brain iron accumulations caused by pilocarpine-induced *status epilepticus* in rats using QSM. Perls staining was performed to visualize the iron deposits as hemosiderin accumulations.

## 2. MATERIALS AND METHODS

### 2.1 Animals

Adult male Sprague-Dawley rats were provided by the Institute of Neurobiology’s animal facility. They were maintained under a 12 h light/dark cycle with access to food and water *ad libitum*. All the rats were acclimatized to the room conditions (20-22°C, 50-60% humidity) for at least one week before any experimental manipulation. The protocol was approved by the Institute of Neurobiology Ethics Animal Care Committee (protocol 105A). The procedures were performed in accordance with the Mexican Federal regulations for animal experimentation (NOM-602-ZOO-1999) and ARRIVE guidelines (Kilkenny et al., 2012). The number of animals was kept as small as possible.

### 2.2 Intracerebral injection of iron chloride

Five healthy rats (60 days old) were used to evaluate the sensitivity of our QSM protocol for detecting brain iron deposits. Animals were anesthetized with an intraperitoneal injection of ketamine/xylazine mixture (70 and 10 mg/kg, respectively) and placed in a stereotactic frame. An incision was made midline along the scalp, and the skull was exposed using sterile surgical techniques. Based on the Compact Paxinos and Watson atlas 6^th^ edition (2009), a small hole was drilled into the skull at the following coordinates: anteroposterior - 5.6 mm, lateral -5.5 mm, ventral from skull surface 7.4 mm. Then, a 5 µl Hamilton syringe was adapted to the stereotactic frame arm and gradually lowered into the right ventral hippocampus to infuse 2 µl of iron chloride (FeCl_3_) solution or vehicle (sterile 0.1M phosphate-buffered saline [PBS]; Sigma-Aldrich) at a rate of 0.1 µl/min using a WPI Microsyringe Pump Controller. Four rats received different concentrations of FeCl_3_ solution (0.1, 1, 10, and 100 mM); the fifth rat received 0.1M PBS. The wound was sutured, and the animals were transferred back to their cages. All the animals were scanned and perfused the next day. The iron deposits were detected by Perls staining (see below).

### 2.3 *In vivo* MRI acquisitions

The acquisition protocol was carried out at the National Laboratory of Magnetic Resonance Imaging using a 7-T Bruker animal scanner interfaced to a Paravision 7.0 console. *In vivo* QSM imaging was acquired using a transmit-receive volume coil with an inner diameter of 72 mm.

Animals were anesthetized with a 5% sevoflurane/oxygen mix. This concentration was reduced by 40% to maintain anesthesia during image acquisition. The respiration rate was monitored and kept at 40-50 breaths per minute. Body temperature was maintained using a warm-water circulation system. Inhaled anesthesia was suspended upon termination of the imaging session. Once the rats were fully recovered, they were transferred back to their polypropylene cages.

Data sets for QSM were acquired using a three-dimensional multi-echo gradient sequence with the following parameters: TR = 47.2 ms, TE1 = 1.942 ms, echo spacing = 2.1 ms, 8 echoes, flip angle = 16.5°, spatial resolution = 200 x 200 x 500 mm ^3^. The scan time was 26 minutes.

### 2.4 Image data processing and analysis

The initial processing was done using brkraw version 0.3.11 (https://brkraw.github.io), MRtrix 3.0, and FSL. Here, raw data were converted to magnitude and phase maps in NifTI format; cerebral maps were also generated in this format (Fig. 1A). The resulting maps were processed with SEPIA version 1.2.1 (https://sepia-documentation.readthedocs.io) and run in MATLAB 2023a (MathWorks, Natick, MA). Then, we followed the next steps to create the susceptibility maps: 1) combination of multiple echoes using the tool Optimum Weights (Schweser et al., 2017); 2) phase unwrapping using the Laplacian method implemented in MEDI toolbox (Zhou et al., 2014); 3) background field removal with the VSHARP method (Li et al., 2011); and 4) creation of QSMs with the STAR-QSM method (Wei et al., 2015). Finally, the maps were exported to NifTI (Fig. 1B).

**Figure 1.**
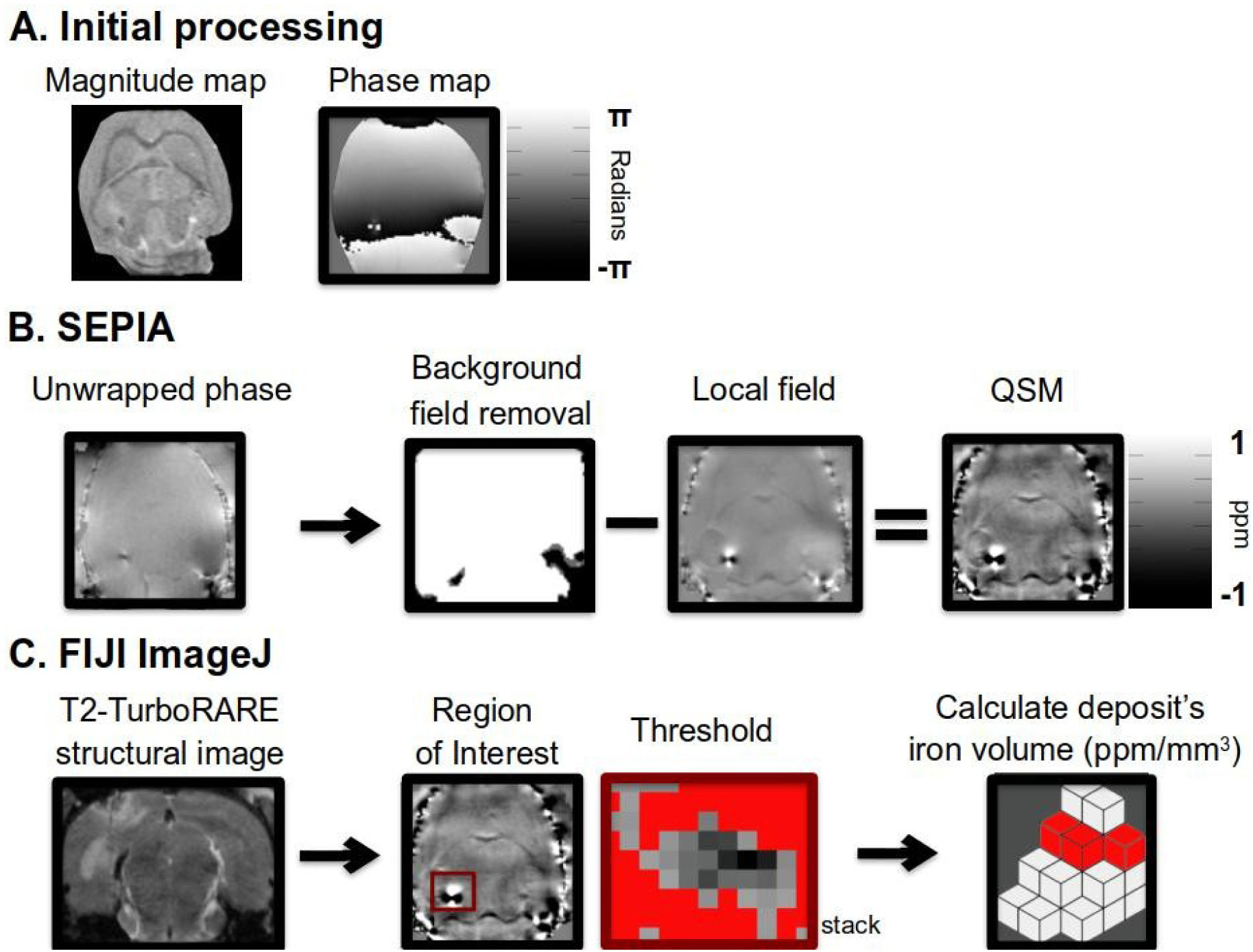
Image processing and analysis. The initial processing consisted of converting the raw data into magnitude and phase maps (A). SEPIA was used as the quantitative susceptibility mapping (QSM) pipeline analysis tool (B); FIJI ImageJ was used for the quantitative analysis of iron deposits identified in the QSM images (C). The red box delineates the zone where the iron deposit was detected (dark hole with a surrounding hyperintensity). The volume of each accumulation was measured (ppm/mm^3^) (C).

MRtrix 3.0 with MRView visual interface was used to detect iron deposits (Tournier et al., 2019). Here, we applied a filter on the QSM maps to easily identify only the paramagnetic intensities (>0.1 ppm). For a precise anatomical localization of the iron deposits, we used the T2-weighted TurboRARE structural images.

FIJI ImageJ was used for the quantitative analysis of the iron deposits identified in the QSM images. Each accumulation was treated as a region of interest, and duplicated in a sub-stack to cover its entire surface in three dimensions. Then, a threshold was applied to evaluate only paramagnetic intensities; the area and intensity of each deposit were measured on each slice. The size of the deposits was calculated by multiplying the paramagnetic area of each slice (mm^2^) by the slice thickness (0.5 mm). The resulting value (mm^3^) of each slice, when added, provides an approximate value of the deposit’s volume. Finally, by multiplying this volume by the mean voxel value, we obtained the deposit’s iron mass (ppm/mm^3^) (Fig. 1C).

### 2.5 Pilocarpine-induced *status epilepticus*

Once we demonstrated that iron deposits can be detected by our QSM protocol, we proceeded to the next step. Nineteen animals were habituated to handling by receiving an injection of saline solution (0.9% NaCl, 1 ml/kg, ip) every day for four days. Then, the animals underwent the protocol described in Luna-Munguia et al. (2017). Briefly, fifteen rats (56 days old) were injected with pilocarpine hydrochloride (340 mg/kg ip; Sigma-Aldrich) 20 min after receiving atropine sulfate (5 mg/kg ip; Sigma-Aldrich). Five to ten minutes after administering the pro-convulsive, the animals exhibited head nodding and myoclonic jerks that evolved into recurrent generalized seizures within 40 to 45 min. Rats that did not develop *status epilepticus* within this time frame, received an additional dose of pilocarpine hydrochloride (170 mg/kg ip). Diazepam (10 mg/kg ip; PiSA) was injected after 90 min of *status epilepticus*. Control animals (n=4) received a single injection of atropine sulfate (5 mg/kg ip; Sigma-Aldrich) 20 min prior to vehicle (0.9% saline solution ip); diazepam (10 mg/kg ip) was injected 2 h after the saline injection. A total of 5 rats died during the *status epilepticus*.

Control (n=4) and *status epilepticus*-induced rats (n=10) were scanned at five time points: prior to *status epilepticus* (50 days old) and 1, 7, 14, and 21 days post-induction. Animals were perfused the day after the last scan. The iron deposits were detected by Perls and diaminobenzidine (DAB) staining (see below).

### 2.6 Brain extraction and histology

Rats were overdosed using an intraperitoneal injection of sodium pentobarbital (Pet’s Pharma). Then, they were intracardially perfused with 0.9% NaCl solution followed by a 4% paraformaldehyde (PFA) solution (pH 7.4). Brains were removed from the skull and post-fixed in fresh 4% PFA solution for 24 h at 4 °C. Then, they were immersed in 20% and 30% sucrose solutions at 4 °C. Each specimen was carefully frozen using dry ice and stored at - 72 °C until slicing.

The stereotactically injected iron was detected by Perls staining. The protocol was partially based on Akyuz et al. (2021). Briefly, brains were sectioned in the coronal plane (40 µm) using a Leica Biosystems 3050S cryostat. The series of sections were preserved in PBS at 4 °C until the mounting and staining day. The 40 µm-thick slices were incubated for 15 min in an equal-volume solution containing 0.2% C₆FeK₄N₆ and 0.2% HCl, both dissolved in deionized water. The sections were counterstained with 0.02% safranine solution. Finally, the sections were mounted on slides and left to dry overnight at room temperature. Images were taken using an AmScope trinocular microscope fitted with a digital camera.

The iron deposits, visualized as brain hemosiderin accumulations in the pilocarpine-injected rats, were detected based on Aggarwal et al. (2018). Here, we enhanced the Perls reaction using DAB. The brain slices were mounted on slides and rinsed three times in PBS. The slides were submerged for 15 min in an equal-volume solution containing 0.5% C₆FeK₄N₆ and 0.5% HCl. Another PBS wash was done, and the slides were incubated for 10 min in DAB solution (0.5 mg/ml dissolved in 0.1M PBS). The DAB reaction was triggered with 0.03% H_2_O_2_ for 15 min. Finally, the slides were washed several times with PBS, dried, and cover-slipped. Images were taken using an AmScope trinocular microscope fitted with a digital camera.

### 2.7 Statistical analysis

The volume of iron deposits was evaluated over time in each brain region where they were detected. Values are expressed as mean ± S.E.M. Results were analyzed by a one-way ANOVA, and a *post hoc* Tukey multiple comparisons test was conducted. In all statistical comparisons, significance was assumed at the level of *p* < 0.05.

## 3. RESULTS

### 3.1 QSM as an effective tool for visualizing iron deposits *in vivo*

We injected four different FeCl_3_ solutions (0.1 to 100 mM) into the ventral hippocampus of four rats as a first attempt to validate our QSM protocol as an effective technique to detect cerebral iron deposits *in vivo*. The animal receiving the highest dose of FeCl_3_ (100 mM) had a couple of generalized seizures 60 minutes after surgery. The acquired QSM images showed evident dark holes with a surrounding hyperintensity in the exact place where the FeCl_3_ solutions were injected. The size of the holes increased according to the concentration of the injected FeCl_3_ solutions (Fig. 2A). To obtain a precise anatomical localization of the iron deposits, we used the T2-weighted TurboRARE structural images. The presence of iron was evident in each of the four injected rats (Fig. 2B). Additionally, the Perls-stained sections revealed considerable damage to the ventral hippocampus of the animal injected with the highest concentration of FeCl_3_ (data not shown). In the other three rats, an intense blue spot was observed at the site of injection. Figure 2C shows the Perls staining section of the animal that received the intrahippocampal injection of FeCl_3_ 10 mM. Neither imaging nor histological effects were observed in the PBS-injected rat (data not shown).

**Figure 2.**
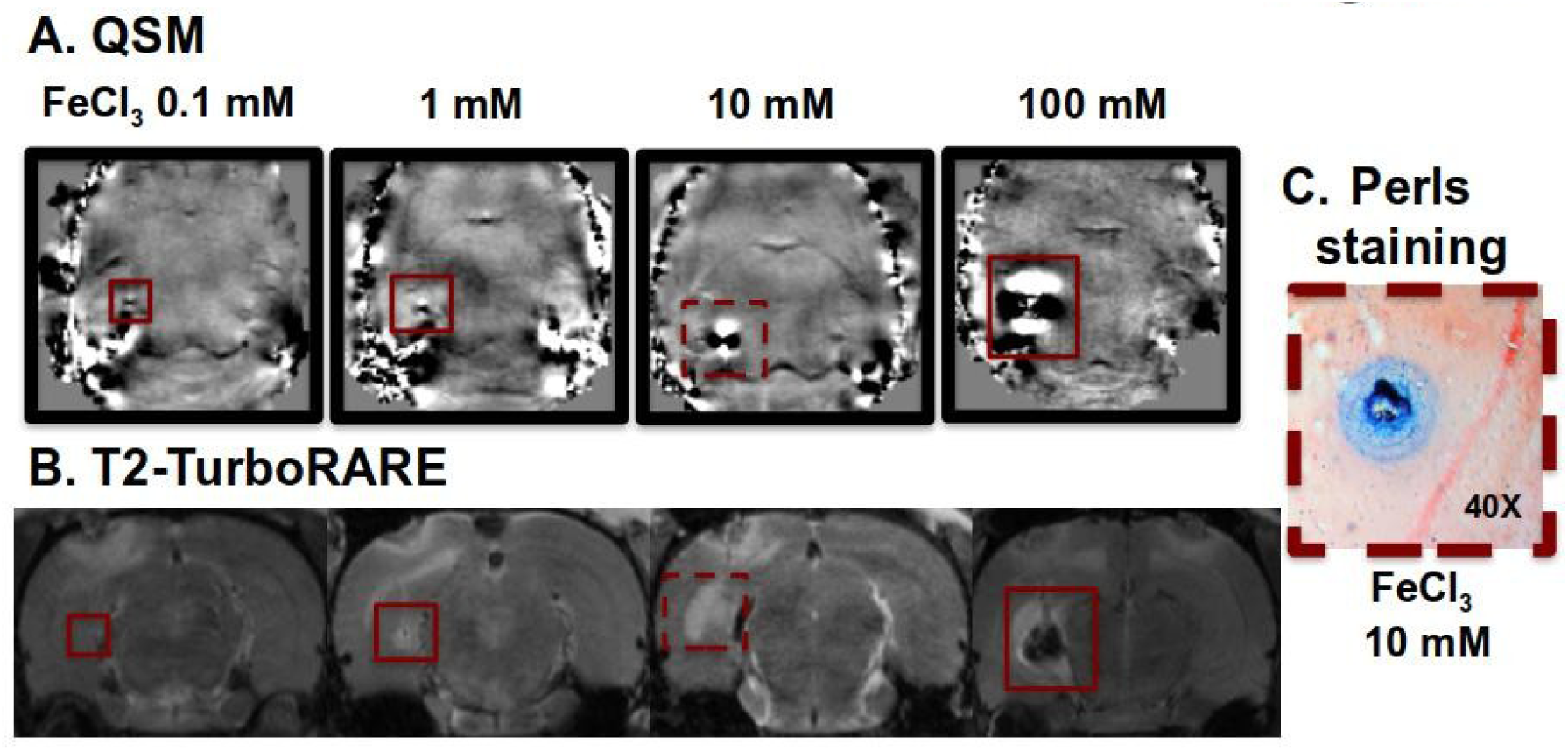
QSM and Perls staining to visualize the injected FeCl_3_ solutions. To validate our QSM protocol, four different concentrations (0.1, 1, 10, and 100 mM) were injected into the ventral hippocampus of four rats. The red boxes show the locations of iron deposits detected *in vivo* (dark holes with a surrounding hyperintensity that increased according to the injected concentration) (A). The coronal view of the T2-weighted TurboRARE structural images was used for a precise anatomical localization of the iron deposits (B). Perls staining confirmed the place of injection. The blue spot correlates with the dark hole-hyperintensity seen in QSM. Here, we show only the picture of the rat that was injected with the 10 mM FeCl_3_ solution (C).

### 3.2 Longitudinal changes in cerebral iron accumulation after pilocarpine-induced *status epilepticus*

QSM images obtained during the basal scan did not show any cerebral iron accumulation (Fig. 3A). Interestingly, all animals had iron deposits (unilateral or bilateral) in the first scan post-*status epilepticus* induction. The accumulations were detected in specific brain regions, such as the caudate putamen, dorsal hippocampus, thalamus, and ventral hippocampus (Fig. 3B). Some animals also showed them in the primary somatosensory cortex (data not shown). When evaluating the size of the deposits during the following weeks, we observed that those localized in dorsal hippocampus, thalamus, and ventral hippocampus increased (Figs. 3C to 3E and 4A), achieving significance only in the ventral hippocampus on the third and fourth scan post-*status epilepticus* induction (\**p* < 0.05 and \*\**p* < 0.01, respectively) (Fig. 4B). The reverse effect occurred in caudate putamen (Fig 4B). The observation of Perls + DAB-stained sections confirmed the abnormal presence of the iron deposits detected by QSM (Fig. 4C).

**Figure 3.**
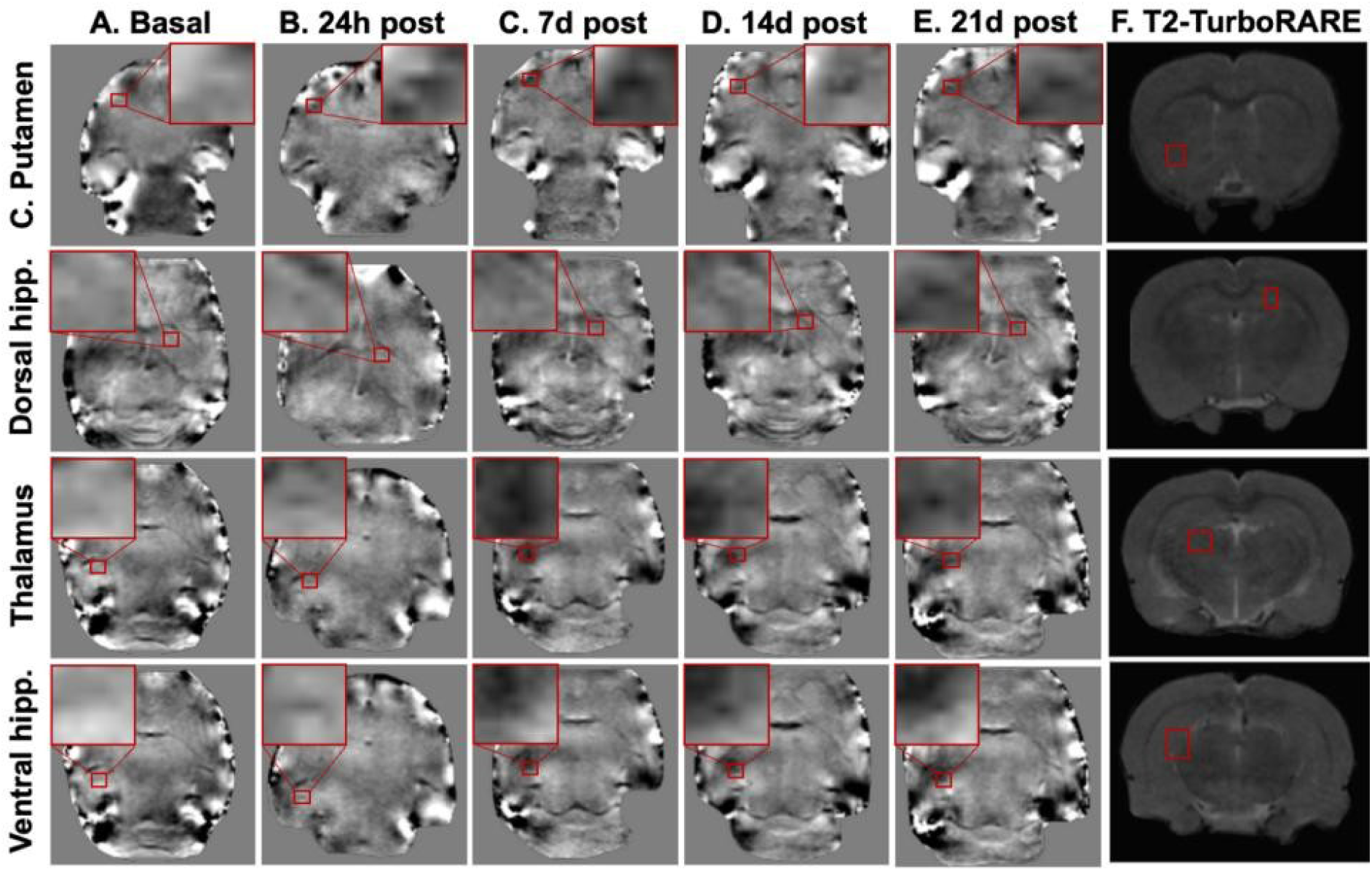
QSM to detect cerebral iron deposits. None of the animals had cerebral iron deposits during the basal scans (A). However, all animals submitted to pilocarpine-induced *status epilepticus* showed iron accumulations in the caudate putamen (C. Putamen), dorsal hippocampus (hipp.), thalamus, and ventral hippocampus (hipp.) 24 h post-induction (B). The non-invasive imaging technique allowed us to evaluate the dynamics of the iron deposits weekly (C-E). The small red boxes show the exact place where these accumulations were detected *in vivo*. The big red boxes show a zoom in of these sites. The coronal view of the T2-weighted TurboRARE structural images was used for a precise anatomical localization of the iron deposits (F).

**Figure 4.**
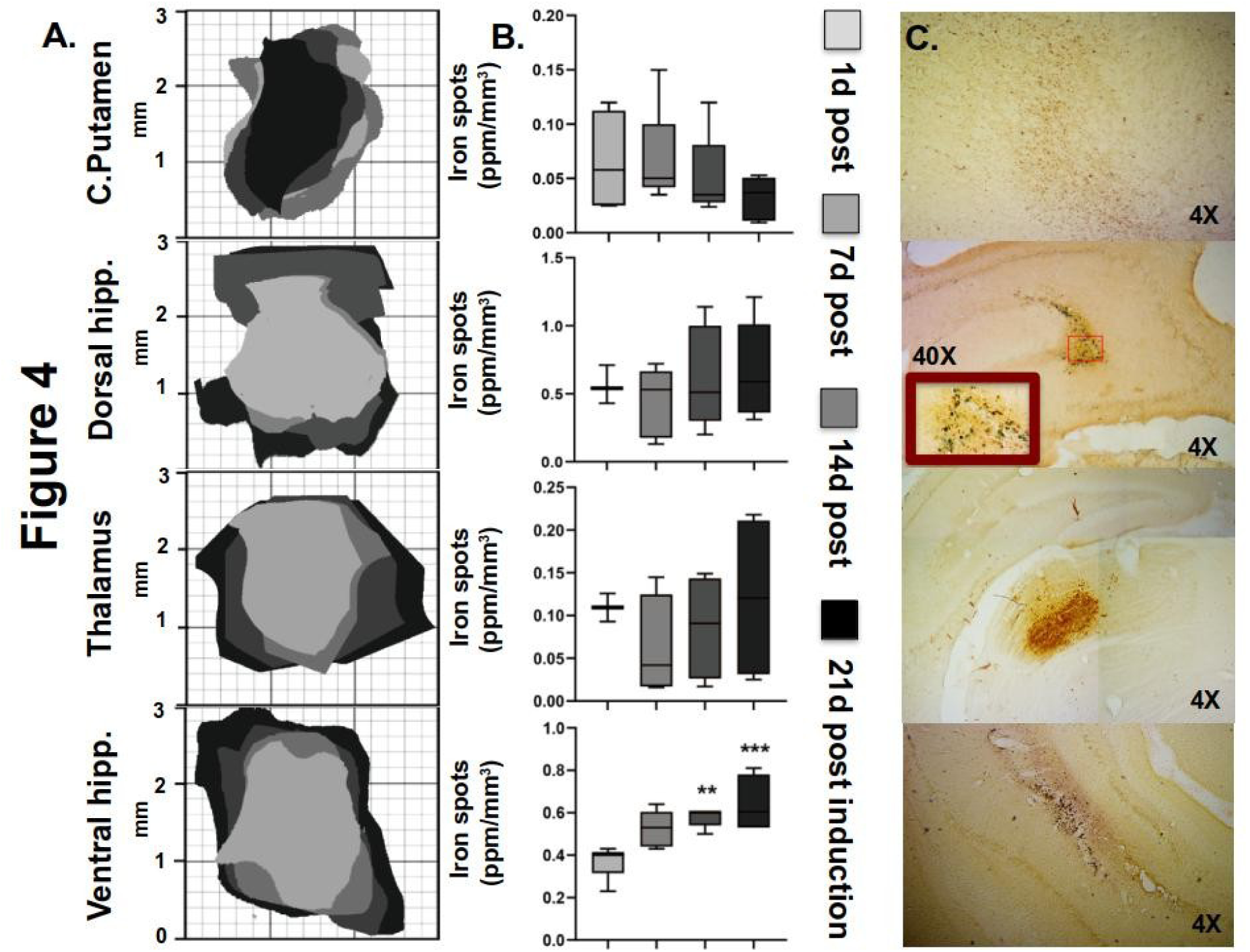
Temporal changes in the cerebral iron deposits. The size of the deposits was evaluated at four different time points (1, 7, 14, and 21 days post-*status epilepticus* induction) in the caudate putamen (C. Putamen), dorsal hippocampus (hipp.), thalamus, and ventral hippocampus (hipp.) (A). Significant differences were only observed in the ventral hippocampus during the third and fourth scans. In contrast to the other three structures, the size of the caudate putamen deposits tended to decrease over time. Data were analyzed by a one-way ANOVA followed by a post hoc Tukey test. Graphs represent the mean ± SEM. \**p* < 0.05; \*\**p* < 0.01 (B). The Perls + DAB-stained sections confirmed the presence of the iron deposits detected by QSM (C).

## 4. DISCUSSION

Iron accumulation and increased ferroptosis markers (ferritin, 4-HNE, MDA) have been described in the epileptic focus resected from patients with refractory temporal lobe epilepsy (Zimmer et al., 2021). Recent studies in animal models of epilepsy showed similar results (Chen et al., 2022, 2025; Wang et al., 2024a). To date, only Aggarwal et al. (2018) have been able to detect iron accumulations in *ex vivo* samples by using other techniques such as QSM. Other reports have even described that treatments focused on reducing ferroptosis clearly reduced seizure severity (Chen et al., 2022; Wang et al., 2022; Wang et al., 2024b; Xie et al., 2022). Based on these references, it seems that epileptic seizures can stimulate ferroptosis, but it is still unknown how iron deposits can induce epilepsy. In this study, we provide evidence supporting the use of QSM to detect brain iron accumulations in a rat model of induced *status epilepticus*. Moreover, this non-invasive technique allowed us to monitor these accumulations longitudinally.

Recently, Auzmendi and Lazarowski (2025) suggested that ferroptosis is not only a seizure-induced consequence but also a cause of epilepsy development; a position mainly based on previous clinical and experimental studies (Hammond et al., 1980; Ikeda, 2001; Wang et al., 2023). To evaluate this hypothesis, we used the pilocarpine-induced *status epilepticus* model. The animals were scanned at four different time points after the induction. From the first QSM scan, we detected iron deposits in the hippocampus, caudate putamen, frontal cortex, and piriform cortex. These are brain regions in which blood brain barrier integrity is immediately compromised after a severe cerebral insult (*status epilepticus*) (Mendes et al., 2019). Therefore, we suggest that the observed iron accumulations in our animals could result from the *status epilepticus* and may predispose to future spontaneous recurrent seizures. This idea is consistent with a recent study in a *status epilepticus* model, in which rats pretreated with CoQ10 and valproic acid exhibited an evident decrease in neuronal death and ferroptosis markers (Fikry et al., 2024). Moreover, consistent with the apparent proconvulsant role of iron, diverse clinical studies have reported that complete removal of hemosiderin during cavernoma surgery is crucial for achieving seizure freedom in patients (Kitaura et al., 2021; Schuss et al., 2020; Zanello et al., 2019). Ferroptosis is therefore now considered a new target in the treatment of epilepsy and also impacts the prevention of sudden unexpected death (Jin et al., 2023; Moscovicz et al., 2024).

Our results clearly show cerebral iron deposits; however, we do not know their origins. Diverse experimental studies have reported brain microbleeds as a consequence of *status epilepticus* induction, suggesting that this phenomenon can provide the required iron to induce hemosiderin formation via hemoglobin degradation (Biagini, et al., 2008; Gualtieri et al., 2012; Kyriatzis et al., 2024). We do not dismiss the possibility that some of the iron originates this way, yet the progressive accumulation into deposits suggests that more complex processes are involved. *In vitro* experiments have shown that lipopolysaccharide (LPS)-treated microglia can alter neuronal iron homeostasis by increasing the synthesis of interleukin-6 (IL-6) and tumor necrosis factor alpha (TNF-α); events that lead to an increased neuronal expression of divalent metal transporter 1 (DMT1), ferritin heavy chain 1 (FTH1), and transferrin receptor 1 (TfR1) (Pandur et al., 2018). Similar results were described by Urrutia and colleagues when using primary cultures of microglia, astrocytes, and neurons treated with IL-6, TNF-α, or LPS (Urrutia et al., 2013). Recent bioinformatic studies have also shown a strong immunological component in the relationship between ferroptosis and epilepsy, suggesting some ferroptosis-related genes as potential markers (TLR4, HIF-1A, and HMOX1) (Chen et al., 2023b; Li et al., 2025; Xu et al., 2023).

As previously mentioned, cerebral iron deposits observed in our animals could be associated with a disruption of the blood brain barrier during the ictal and interictal periods (Breuer et al., 2017; van Vliet et al., 2007). Once the blood brain barrier is damaged, it starts the extravasation of serum proteins and other blood-derived factors into the brain parenchyma, facilitating the development of spontaneous recurrent seizures, neuronal loss, and astrogliosis (Cacheaux et al., 2009; Tomkins et al., 2007). Moreover, barrier leakage also allows iron to enter from other sources, such as iron bound to heme secondary to hemorrhage, iron bound to transferrin or ferritin, and free plasma iron. Under inflammatory conditions, ferritin levels significantly increase as a consequence of free iron sequestration to mitigate oxidative stress. However, when iron bound to ferritin crosses the compromised blood brain barrier, it can induce a localized iron overload, resulting in neuronal injury and ferroptosis (Petzold et al., 2011). In line with this, Garcia-Yebenes and colleagues (2012) reported that high levels of iron (measured as serum ferritin) are associated with a worse outcome after stroke induction in mice. Yuan and colleagues (2021) conducted a two-sample Mendelian randomization study and suggested that elevated levels of iron, ferritin, and transferrin in serum may be associated with an increased risk of developing epilepsy.

Perls and DAB staining performed in this study show some fine dots that may correspond to extracellular iron deposits united by a ferritin core. In tissues, ferritin is an intracellular protein able to store nearly 4,500 iron atoms per molecule, releasing them when necessary via ferritinophagy. *In vitro* studies have shown that ferritin contributes to sustained iron homeostasis; it is released by exocytosis due to increased extracellular iron levels (Chiou & Connor, 2018; Yu et al., 2023). According to one report, mice with reduced mitochondrial ferritin exhibit neuronal ferroptosis and hippocampal epileptic activity, which are attributed to increased intracellular free iron ions and a decreased function of iron-sequestering proteins (Song et al., 2024). On the other hand, another explanation for the dotted marks we observed may be related to iron, microglia, and pro-inflammatory processes. Recent studies report the involvement of microglia in ferroptosis induced by stroke, ischemia/reperfusion, and amyotrophic lateral sclerosis (Shi et al., 2024b; Swanson et al., 2023; Wang et al., 2024c). In this regard, *in vitro* experiments by Urrutia and colleagues (2013) showed that microglial stimulation (with IL-6, TNF-α, or LPS) facilitates hepcidin secretion and DMT1 expression, promoting detrimental iron accumulation in the cells. Other authors recently described that some anti-inflammatory treatments promote physiological iron storage by decreasing ferritin heavy chain expression and increasing ferritin light chain expression (Fei et al., 2024; Wang et al., 2024c).

Epileptogenesis is a complex multifactorial pathologic process that precedes spontaneous recurrent seizures and subsequent neuronal damage (Scheffer et al., 2017). In this context, brain iron accumulation has been considered not only a predisposing factor but also a key pathogenic brain alteration involved in comorbidities such as depression and cognitive impairment (Duan et al., 2022; Hooper et al., 2018; Keezer et al., 2016; Kiersnowski et al., 2023; O’Toole et al., 2014; Sankar and Mazarati, 2010; Sun et al., 2021; Wulsin et al., 2016). When considering iron as a factor involved in the onset, development, and progression of epilepsy, we cannot discard the role of hemosiderin. A recent study described that even in thalamic tumors without cortical involvement, seizure occurrence is not uncommon (17%); therefore, the thalamus may have a direct role in epileptogenesis (Choon et al., 2023).

*In vitro* studies using ferromagnetic nanoparticles demonstrated that intracellular non-heme iron accumulation is safe at certain concentrations (approximately 50 µg/ml) (Lee et al., 2020). On the other hand, higher concentrations promote oxidative stress and cell death (Hohnholt et al., 2013; Singh et al., 2012). Unlike iron overload due to iron storage and transport proteins, nanoparticles have coatings that facilitate their entry into the cell (ranging from diffusion to pinocytosis) and prevent extracellular accumulation (Wei et al., 2021). Once inside the cells, nanoparticles are stored in vesicles and degraded in a similar way to ferritinophagy. This suggests that cells have a buffer capacity to store a certain excess of iron without impacting viability.

## 5. LIMITATIONS

It is important to mention some considerations about our study. First, QSM methods are affected by residual background field contributions and artifacts; specifically, near air-tissue interfaces. While these effects are minimized in deep brain structures, which comprise the majority of our findings, it is complicated to detect the cortical iron deposits. Overcoming this challenge will add valuable information to our data when evaluating iron dynamics in animal models. Second, noise and the point-dipole problem also complicate the accuracy of the measured susceptibilities (Deistung et al., 2017).

## 6. CONCLUSION

Our study offers compelling evidence of iron deposits after *status epilepticus* induction. Moreover, it proves that QSM is a useful non-invasive technique for detecting these deposits longitudinally. While this study is preclinical in nature, our findings underscore the potential of using QSM for clinical applications where cerebral iron accumulation is suspected (post-traumatic, post-stroke, post-ischemic, post-infection) (Merelli et al., 2021; Ravanfar et al., 2021). Future experiments are crucial to determine the origin of the iron that accumulates in specific brain regions and understand/prevent the vicious cycle that occurs after the insult is triggered (iron deposits seizures new iron deposits).

## Ethics approval and consent

Research was approved by the Institute of Neurobiology Ethics Animal Care Committee (protocol 105A).

## Author’s contribution

FM, AL, LC, JA, and HL-M contributed to the conception, methodology, and design of the study. FM and HL-M performed imaging acquisition and histological procedures. FM and L-VM performed QSM imaging and statistical analyses. LC provided the code and supervised the analysis of iron detection. FM, AL, JA, and HL-M wrote, reviewed, and edited the first manuscript. All authors reviewed the manuscript and approved the submitted version.

## CRediT authorship contribution statement

**Franco Moscovicz:** Data curation, Formal analysis, Investigation, Methodology, Writing – original draft. **Leonardo Vazquez-Morales:** Data curation, Formal analysis, Methodology. **Alberto Lazarowski:** Conceptualization, Writing – original draft. **Luis Concha:** Formal analysis, Data curation, Funding acquisition, Methodology, Resources, Software, Supervision, Validation. **Jeronimo Auzmendi:** Conceptualization, Formal analysis, Methodology, Supervision, Validation, Visualization, Writing – original draft, Writing – review and editing. **Hiram Luna-Munguia:** Conceptualization, Data curation, Formal analysis, Funding acquisition, Investigation, Methodology, Project administration, Resources, Supervision, Validation, Visualization, Writing – review and editing.

## Consent for publication

Not applicable.

## Acknowledgements

We thank Mirelta Regalado, Juan Ortiz-Retana, Ericka de los Rios, Nydia Hernandez-Rios, Lourdes Palma, Moises Mendoza, and Leopoldo Gonzalez-Santos for their technical assistance. We also thank the personnel at the Animal Facility: Martin Garcia, Alejandra Castilla, Maria Carbajo-Mata, and Maria Eugenia Ramos. Jessica Gonzalez-Norris for proof-reading and editing our manuscript. MRI preclinical scanner facilities were provided by the National Laboratory of Magnetic Resonance (LANIREM), which receives support from CONAHCYT and UNAM. Computing infrastructure was partially provided by the National Laboratory for Advanced Scientific Visualization (LAVIS).

## Data availability

All data generated and analyzed during this study will be made available by the authors, without undue reservation.

## Declaration of generative AI use

Authors declare no use of generative AI in the manuscript preparation process.

